# Juvenile Hormone Signalling Underlies the Switchpoint and Differentiation of Soldiers in *Camponotus floridanus*

**DOI:** 10.1101/2025.08.25.672237

**Authors:** Olivia MacMillan, Julia Singer, Sophia Perrakis, Alex Craig, Davina Ntanga, David Qiu, Rajendhran Rajakumar

## Abstract

Multicellular organisms are composed of various cell types derived from cellular determination and differentiation, which carry out specialized functions in the individual. Ant societies are composed of various castes of individuals derived from caste determination and differentiation, which divide specialized tasks in the colony. From cells to societies, the underlying mechanisms orchestrating determination and differentiation at different biological levels are fundamental in generating biological diversity. While past work has described the developmental process of caste determination and differentiation in ants, the endocrinological and molecular basis for these processes in the genus *Camponotus* remains unknown. Here we show that minor worker and soldier development is determined by a Juvenile hormone (JH)-mediated minor-soldier switchpoint in *C. floridanus*. Hormonal treatments identified a sensitivity period when JH can induce bipotential larvae to become soldiers. Induced soldiers phenocopy natural soldiers in size and soldier-specific head-to-body allometry and this is associated with a heterochronic shift in metamorphosis. Furthermore, we molecularly characterize the activity of the JH pathway at the level of synthesis, degradation, reception, and downstream effectors during caste determination and differentiation. Surprisingly, rather than JH synthesis, we found JH reception and JH degradation are the major signalling processes differentially regulated across castes and RNAi of juvenile hormone degradation enzymes generates soldiers. Finally, our data suggests that the hyperdiverse genus *Camponotus* has independently evolved a JH-mediated minor-soldier switchpoint similar to that known in *Pheidole*. More generally, the evolution of plastic hormonal regulation may facilitate the origin of developmental determination and differentiation processes underlying complex adaptive phenotypes.

**Significance Statement:** Ants live in complex societies like a superorganism where individuals divide reproductive and various non-reproductive labors, which are analogous to the array of functions of germ cells and highly differentiated somatic cells in an organism. While there are over 1500 species of the hyperdiverse genus *Camponotus*, a genus whose diversity reflects their complex caste system including minor workers and soldiers, the molecular and endocrinological basis for how these castes develop remains unknown. Here we have identified a juvenile hormone (JH) mediated minor-soldier switchpoint during larval development, and have characterized components of the JH signalling pathway that regulate soldier development. This work enables further understanding of how castes are regulated across ants and social insects more broadly.

## Introduction

Ants are found across almost all continents and terrestrial habitats and have played a major role in ecosystems for millions of years (1, 2). This ecological success has been attributed to the origin of eusociality and the complex caste systems present within ant colonies (2). Eusociality in ants is the reproductive division of labour where there are winged reproductive individuals (males and queens) that perform reproductive tasks, and wingless sterile individuals (workers) that perform non-reproductive tasks (1). Several ant species have independently evolved the ability to further differentiate their worker caste into minors that nurse and forage for food, and majors (also known as soldiers) that defend the colony and aid in food processing (1, 3, 4). To facilitate optimality of these different tasks, there are head-to-body allometric differences between the small-headed minors and big-headed soldiers (4–6). Although, caste differences and individual responsibilities vary greatly, females in the colony are highly related and therefore the female castes and variation within are not due to genetic variation alone. Rather, the action of hormonal and epigenetic mechanisms provide a means for phenotypic variation to emerge from a similar genetic background in response to environmental variation acting on development (1, 4, 7–9).

Developmental plasticity is the ability for a genotype to give rise to phenotypic variation as a result of environmental variation acting on development (10, 11). In ants, variation of several environmental factors are known to influence whether an individual develops into a minor or a soldier including temperature (12) and nutrition (12–15). Mechanisms that regulate Gene-by-environment (GxE) interactions are imperative for caste determination, differentiation, and growth in ants and include hormonal and epigenetic mechanisms (5, 7, 16–18).

Previous work investigating GxE interactions in the context of hormones for queen-worker and minor-soldier caste determination in ants has focused on juvenile hormone (JH). JH increases at critical developmental time points to mediate ant caste determination (16, 18–20). The first JH-mediated switchpoint controls queen-worker caste determination and the developmental timing for when this occurs can vary greatly across species (1, 20). In the species *Pheidole pallidula* (16) the JH-mediated queen-worker switchpoint occurs during embryonic stage. In contrast, the queen-worker switchpoint is during the 3^rd^ (final) instar in *Myrmica rubra*, (21), and during the 3^rd^-4^th^ instars in *Harpegnathos saltator* (22).

Nearly half a century ago, it was first demonstrated in the hyperdiverse (over 1000 species) ant genus *Pheidole* that soldier development is determined by a second JH-mediated switchpoint (Fig. 1*B*) in the last larval instar (18). The comparably hyperdiverse ant genus *Camponotus* is separated from *Pheidole* by over 100 million years of evolution (23), and has independently evolved a soldier caste (Fig. 1*A*). However, nothing is known about the hormonal mechanisms influencing minor-soldier caste-specific development in *Camponotus* (Fig. 1*C*). Here we sought to characterize at both the molecular and hormonal levels how soldiers are determined and differentiate in *Camponotus*. Using *Camponotus floridanus* an emerging ant model for behavioural, genomic, epigenetic and social complexity studies (7, 24–29), we investigated the impact of JH on adult sizing, head-to-body allometry, and developmental timing as well as assessed, at the molecular level, which JH pathway components were critical in influencing caste-specific development.

**Figure 1.**
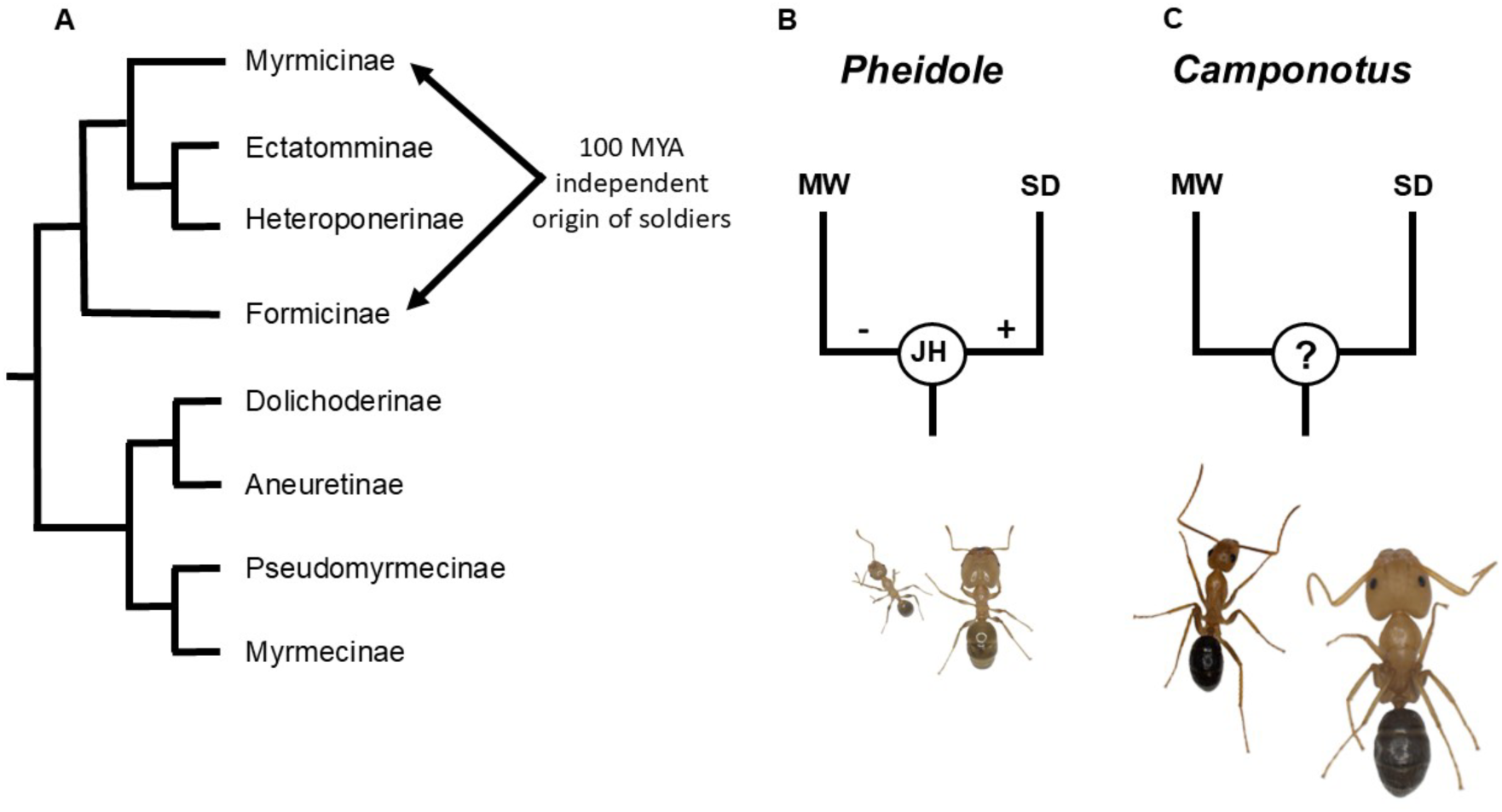
Juvenile hormone pathway and the independent evolution of soldier development in ants. (A) A simplified ant phylogeny based on Romiguier et al. (2022), demonstrates the evolutionary separation of two hyperdiverse genus *Pheidole* and *Camponotus* spanning more than100 million years. (B) Pheidole worker castes are mediated by a JH caste-specific switchpoint. (C) Whether there is a minor-soldier switchpoint in *Camponotus* and if it is regulated by JH remains unknown. Adults in (B) are a minor and soldier of *Pheidole pallidula* and (C) *Camponotus floridanus*.

## Results

### JH regulates caste-specific sizing and developmental timing in *C. floridanus*

Separated by 100 million years, we wanted to determine if there is a JH-mediated minor-soldier switchpoint in *Camponotus* similar to that found in *Pheidole*. Based on past larval instar characterization in *C. floridanus,* the majority of larval growth happens in the 4^th^ and final instar (7), and therefore we predicted that the JH-mediated minor-soldier switchpoint would occur during that stage or just prior. To explore this, we conducted hormonal manipulations at three candidate developmental periods including the 3^rd^ instar, the beginning of the 4^th^ instar, and the middle of the 4^th^ instar. The larvae from each stage were treated with a synthetic analogue of JH (methoprene) and left to develop to adulthood for head width and scape length measurements; scape length being a known proxy for body size in *Camponotus* (30) and *C. floridanus* specifically (7). For the 3^rd^ instar hormonal treatments, the head width (16.4% increase, Fig. S1*D*, p<0.0001) and scape length (7.8% increase, Fig. S1*A*, p<0.0001) were significantly larger for methoprene-treated individuals compared to controls. Minor workers and soldiers differ in their head-to-body allometry and therefore using a linear regression we compared the slope of methoprene-treated individuals to the control group and found that while there was no significant difference between slopes (p=0.1708), there was a significant difference between y-intercepts (p=0.0041) (Fig. 2*A*). For the early 4^th^ instar treatments, the head width (23.4% increase, Fig. S1*E*, p<0.0001) and scape length (7.6% increase, Fig. S1*B*, p= p<0.0001) were significantly larger compared to controls, and the slopes were significantly different (Fig. 2*B*, p<0.0001). Finally, for the middle 4^th^ instar treatments, the head width (33.9% increase, Fig. S1*F*, p<0.0001) and scape length (9.3% increase, Fig. S1*C*, p<0.0001) were significantly larger, and the slopes were significantly different (Fig. 2*C*, p<0.0001). Together this indicates that methoprene treatments for all groups made the individuals larger relative to their control counterparts and the middle 4^th^ instar was the developmental period most sensitive to juvenile hormone manipulation, in terms of increases in head size, body size, and head-to-body allometry akin to soldiers (Fig. 2*C* and Fig. S1*C* and *F*). In order to better understand the soldier induction potential of these manipulations across the three developmental stages, we compared the mean scape size of the three treatment groups to scape size frequencies in a mature colony of *C. floridanus* previously described (7). While the 3^rd^ instar treatment led to an increase in scape size, only the two 4^th^ instar treatments led to the generation of scape sizes reflecting body sizes comparable to natural soldiers with the middle 4^th^ instar exhibiting the highest similarity (Fig. S1*G*). Altogether, there exist a JH-dependent minor-soldier switchpoint during the 4^th^ instar in *C. floridanus*.

**Figure 2.**
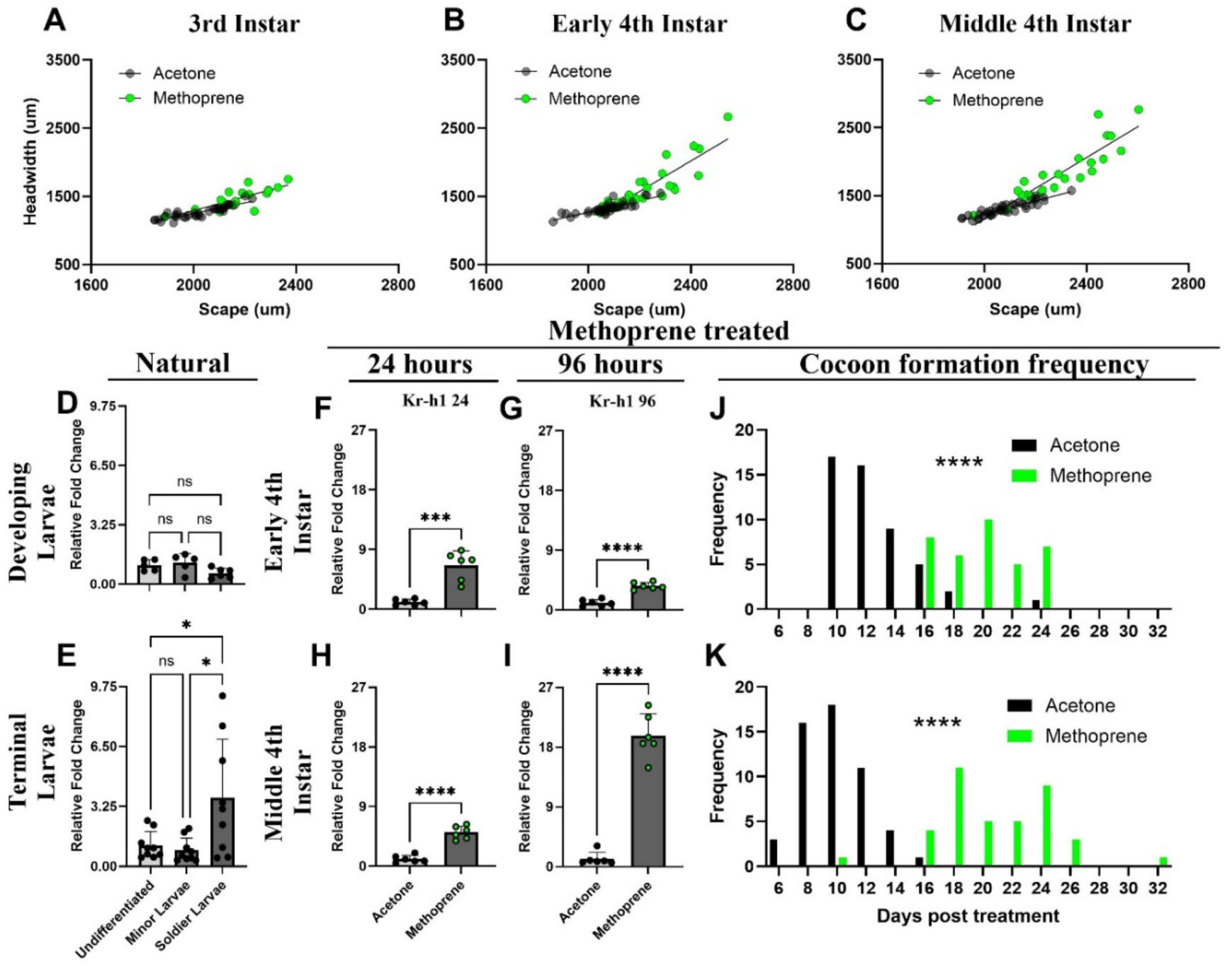
Hormonal manipulations pinpoint the minor-soldier caste switchpoint in *C. floridanus* at molecular and phenotypic levels. (A-C) Allometry of individuals generated after methoprene treatment of 5mg/ml when treated at (A) 3^rd^ instar, (B) early 4^th^ instar, and (C) middle 4^th^ instar. (A) Allometric slopes (p=0.1708) and y-intercepts (p=0.0041) were compared between acetone control and methoprene treatment during the 3^rd^ instar (acetone n=36; methoprene n=18), (B) slopes of 4^th^ instar treatments (p<0.0001) (acetone n=43; methoprene n=26), and (C) slopes of middle 4^th^ instar treatments (p<0.0001) (acetone n=42; methoprene n=27). Simple linear regressions were conducted to determine the statistical significance between acetone and methoprene treatments. (D, E) Relative expression of *Kr-h1* through qPCR determined for (D) developing (p=0.0603 for minor vs soldier) and (E) terminal larva (p=0.0120 for minor versus soldier). The sample size was n=6-9. (F-I) Relative expression differences of *Kr-h1* through qPCR for methoprene-treated early 4^th^ instar larvae versus control at (F) 24 hours post-treatment, (G) 96 hours post-treatment (F; p=0.0001 and G; p<0.0001), and methoprene-treated middle 4^th^ instar larvae versus control at (H) 24 hours and (I) 96 hours post-treatment (H; p<0.0001 and I; p<0.0001). Sample size for qPCR experiments were n=6. Developmental timing in the form of cocoon formation is displayed across (J) early 4^th^ instar (p<0.0001) treated with methoprene (n=26) compared to acetone (n=43), and (K) middle 4^th^ instar (p<0.0001) treated with methoprene (n=27) compared to acetone (n=42). Methoprene treated individuals are represented in green and control samples in black.

Wheeler & Nijhout (1981) demonstrated that *Pheidole* soldier larvae extend their last larval instar compared to developing minor workers, providing an extended period of growth (18). To determine if *C. floridanus* soldier larvae extend their growth period in a similar way, we tracked their developmental timing (cocoon formation and eclosure timing) following the three methoprene treatments (Table S1 and Table S2). The 3^rd^ instar treatments had a 61% (8.25 days) developmental delay in first pupation (Table S1; p=0.0129) and the first individual eclosed with a 16% (6 days) delay in the methoprene-treated group as compared to the control (Table S2; p=0.0751). The methoprene-treated early 4^th^ instar larvae had a 68% (7.5 days) developmental delay in first pupation (Table S1; p<0.0001) and the first individual eclosed with a delay of 32% (11 days) compared to the control (Table S2; p=0.0002). The middle 4^th^ instar-treated larvae had a 161% (11 days) developmental delay in first pupation (Table S1; p<0.0001) and the first individual eclosed with a delay of 51% (16 days) compared to the control (Table S2; p<0.0001). Overall, the pupation frequency was significantly delayed (p<0.0001) in both early (Fig. 2*J*) and middle 4^th^ instar (Fig. 2*K*) treatments, with the middle 4^th^ instar treatment being the most delayed. Therefore, the developmental timing data is consistent with the sizing data suggesting that during the 4^th^ larval instar in *C. floridanus*, there is a JH-mediated switchpoint and that when JH induces soldier development, this instar is extended to enable an increase in growth, similar to that which has evolved in *Pheidole*.

### Molecular signatures of JH signalling during caste determination

Alongside our scaling and developmental timing findings, we wanted to further confirm at the molecular level, the precise timepoint during the 4^th^ larval instar for which soldier induction is most sensitive. *Kr-h1* is an extensively investigated highly conserved transcriptionally-regulated downstream effector of the JH pathway (31, 32). We first looked at *Kr-h1* expression before caste determination, during minor and soldier differentiation, and towards the end of caste-specific differentiation. We found that for *Kr-h1*, there are no caste-specific differences at the beginning of caste differentiation (Fig. 2*D*), yet there is towards the end of caste differentiation, where soldier expression is significantly higher compared to minor workers (Fig. 2*E*). Following this, we did a time course (24-, 48-, 72-, or 96-hours) analysis of JH pathway activity by measuring *Kr-h1* gene expression levels following our hormonal manipulations. We focused on the two 4^th^ instar groups because both resulted in head-to-body allometric differences in slope and extensions of larval developmental timing. Both early 4^th^ instar (Fig. 2*F* and *G*, Fig. S2*A*-*D*) and middle 4^th^ instar (Fig. 2*H* and *I*, Fig. S2*E*-*H*) methoprene treatments led to the upregulation of *Kr-h1*. Furthermore, we found that while the fold change of *Kr-h1* increases rapidly at first and the magnitude of this increase diminishes over time in response to methoprene treatment of early 4^th^ instar larvae (24hr: 7-fold, 48hr: 7-fold, 72hr: 6-fold, 96hr: 4-fold increase; Fig. S2*A*-*D*), larvae treated at the middle 4^th^ instar had *Kr-h1* expression levels that continued to increase in fold change throughout the duration of the 4 timepoints in response to methoprene treatment (24hr: 5-fold, 48hr: 9-fold, 72hr: 17-fold, 96hr: 20-fold increase; Fig. S2*E*-*H*). Collectively, in the context of our JH pathway manipulations, the magnitude and sustained increase in *Kr-h1* expression (Fig. S2*E*-*H*) combined with our allometric (Fig. 2*C*) and developmental timing data (Fig. 2*K* and Table S1 and 2) altogether suggests that larvae are most sensitive to caste determination and soldier induction during the middle of the 4^th^ instar.

### Thermal plasticity of the JH-switchpoint in sizing and soldier production

We next wanted to investigate in the context of GxE interactions, how environmental variation can influence the capacity of JH to induce soldiers. It is known in *Pheidole* that increasing temperature can lead to an increase in soldier production (12). We therefore reared larvae during the minor-soldier switchpoint (middle 4^th^ instar) at 27°C following a methoprene treatment. Raising methoprene-treated individuals at 27°C led to a significant increase in head size (62.3% increase, Figure 3*C*; p<0.0001), scape length (15.6% increase, Fig. 3*B*, p<0.0001), and head-to-body allometry (Fig. 3*A*; p<0.0001) compared to controls. Methoprene delayed first pupation by 146% (Table S3; 9.3 days; p=0.0006) and delays first eclosure by 42% (Table S3; 9.4 days; p=0.0008). Methoprene significantly delays overall pupation frequency (Fig. 3*D*; p<0.0001) and eclosure (Fig. 3*E*; p<0.0001). To further understand how temperature can enhance the soldier induction potential of JH during the minor-soldier switchpoint, we compared the mean scape size of the adults generated from the methoprene treatments and control at 25 °C alongside the methoprene treatment and control at 27°C to scape size frequencies previously described (7). We found that temperature can increase the size of both minors and soldiers and that the combination of both JH and temperature increase led to the production of individuals larger than soldiers typical of a mature colony (Fig. 3*F* and *G*). Altogether, *C. floridanus* larval sizing is developmentally plastic to temperature and JH together with thermal plasticity can act synergistically to induce soldier development.

**Figure 3.**
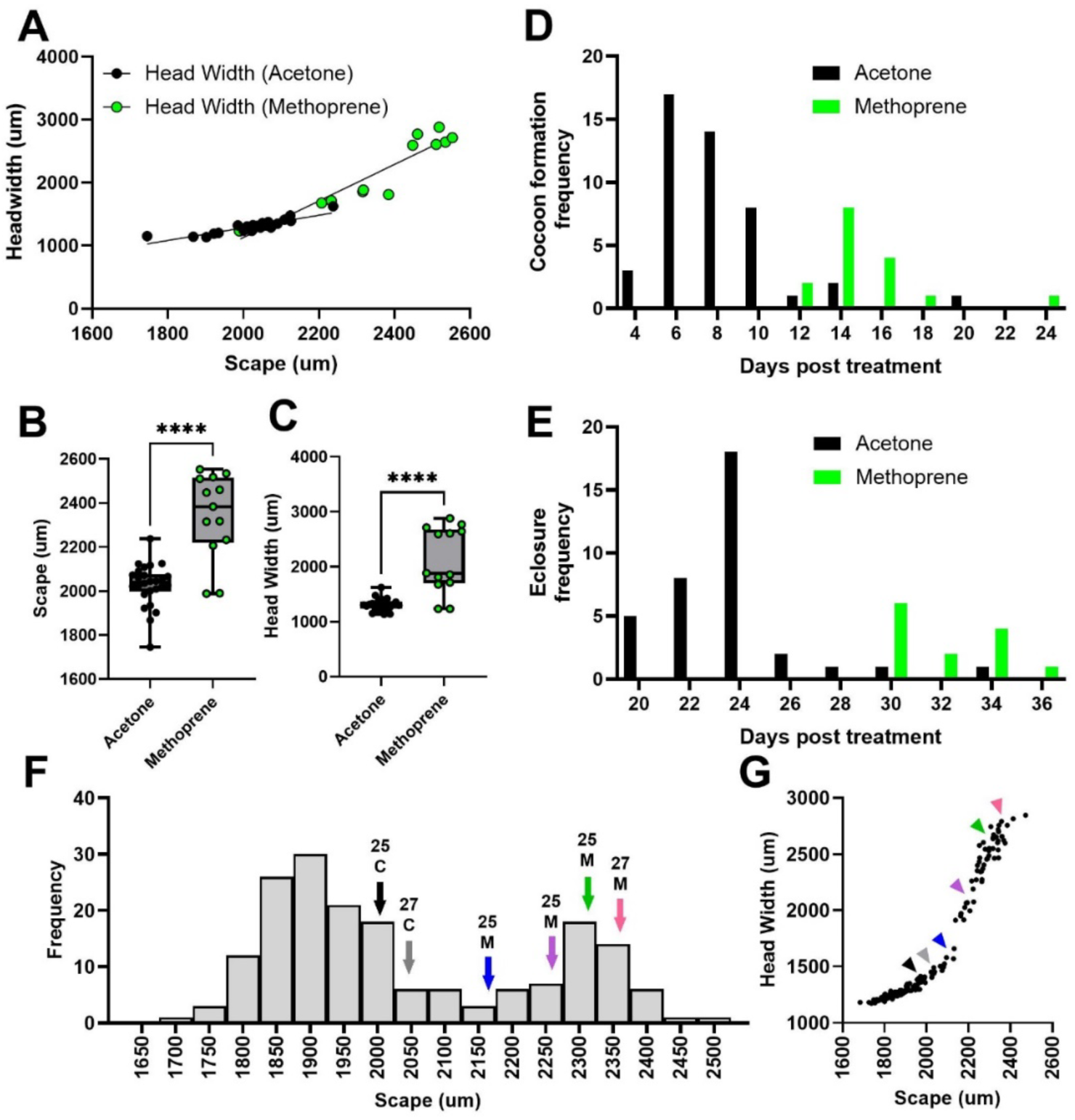
JH and temperature are critical for regulating caste determination and developmental timing. (A) Allometry of individuals generated following a 5mg/ml methoprene treatment and reared at 27°C during the minor-soldier caste switchpoint (middle 4^th^ instar; p<0.0001). (B) Scape lengths (p<0.0001) and (C) head widths (p<0.0001) were compared between methoprene treatment and acetone control. Developmental timing in form of (D) cocoon formation frequency (P<0.0001) and (E) eclosure frequency (p<0.0001) were compared between treatment and control. The sample size for control and treatment are n=26 and 13 respectively. Methoprene treated individuals are represented in green and acetone control is represented in black. (F) Arrows and (G) arrowheads representing the acetone control (black 25°C and grey 27°C) and methoprene treatment of 3^rd^ instar (blue; 25°C), early 4^th^ instar (purple; 25°C), middle 4^th^ instar (green; 25°C), and middle 4^th^ instar (pink; 27°C) demonstrates the scape lengths of treated samples in comparison to scape measurements across a mature *C. floridanus* colony adapted from that previously described (7).

### JH degradation and reception associated with caste determination

Now that we have established the existence and timing of the JH-mediated minor-soldier caste switchpoint in *C. floridanus*, we next wanted to determine how JH signalling is regulated in a caste-specific manner to differentiate soldiers from minor workers. JH signalling is regulated at three levels: (1) synthesis, (2) reception, and (3) degradation. Therefore, we analyzed how these regulatory levels changed across larvae that are undergoing caste-specific differentiation (‘developing’) and larvae that are towards the end of caste-specific differentiation (‘terminal’) for developing minor workers and soldiers. For synthesis, we looked at *JHAMT*, the highly conserved synthesizer of JH in insects (33) and found it did not change between castes developmentally (Fig. 4*A*; p=0.9082) or terminally (Fig. 4*B*; p=0.9773). For reception/sensitivity, there are two highly conserved axes of JH reception at the membrane and nuclear levels. At the nuclear level, similar to *Tribolium*, ants have a single nuclear receptor *Methoprene-tolerant/Germ-cell expressed* (*MET*/*Gce*), which is the pro-ortholog of *MET* and *Gce* of *Drosophila melanogaster* (34, 35) and its co-receptor *Taiman* (*Tai*) (36). At the cell surface level, there are two membrane receptor tyrosine kinases (RTK) *Fibroblast growth factor receptor 1* (*FGFR1*) and *Cadherin 96ca* (*Cad96ca*) (37). We found that while *MET* lacked caste-specific differences developmentally (Figure 4*K*; p=0.3910) and terminally (Fig. 4*L*; p=0.6565), *Tai* is higher in expression in developing and terminal soldiers as compared to minors (Fig. 4*M*; p=0.0095 and *N*; p=0.0335).

**Figure 4.**
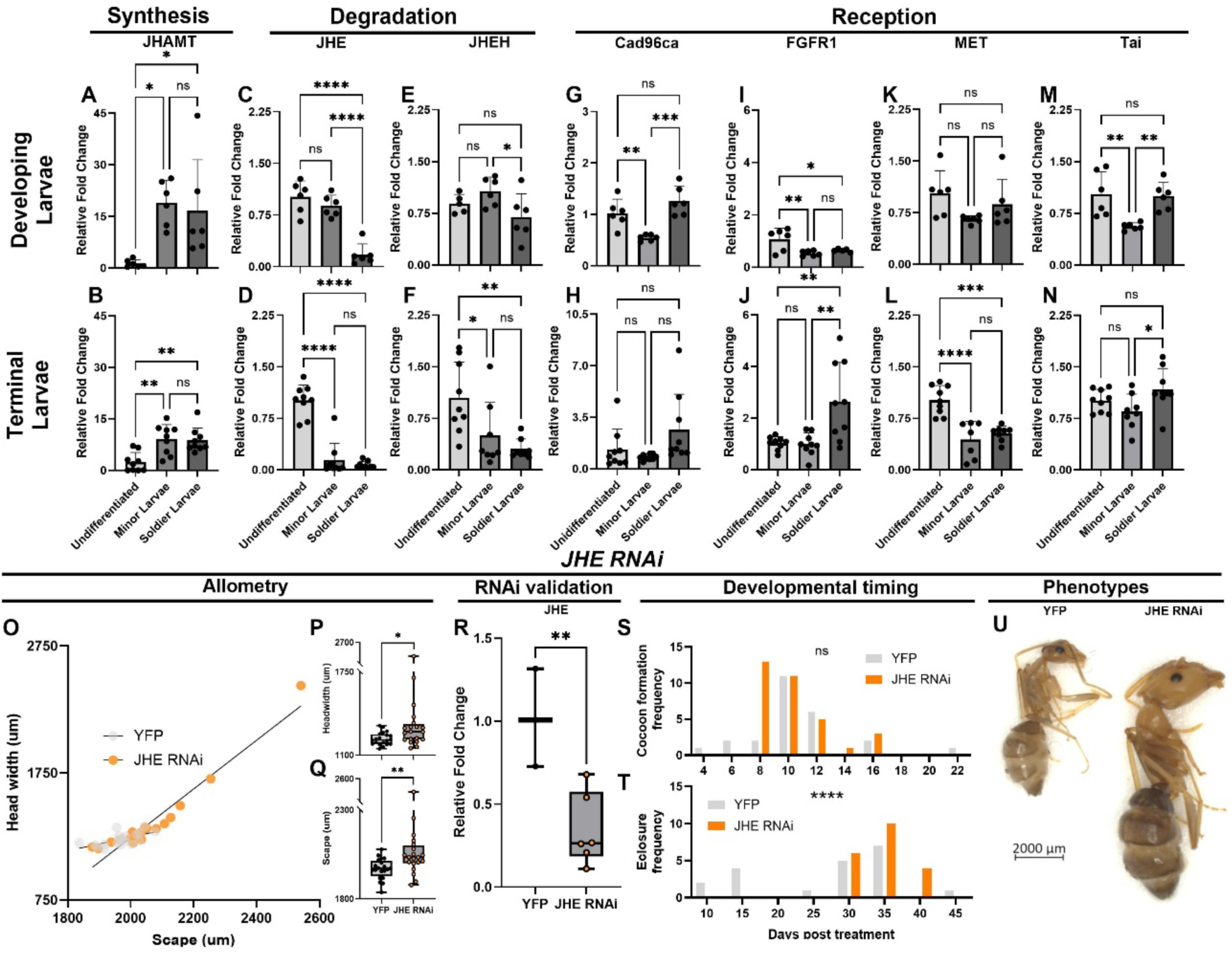
JH signalling during *C. floridanus* soldier development and the role of JH degradation. Relative expression levels of (A, B) *JHAMT*, (C, D) *JHE*, (E, F) *JHEH*, (G, H) *Cad96ca*, and (I, J) *FGFR1*, (K, L) *MET*, (M, N) *Tai*, in *C. floridanus* (A, C, E, G, I, K, M) developing larvae and (B, D, F, H, J, L, N) terminal larvae. Differences between worker and soldier developing larvae (A; p=0.9082, C: p=<0.0001, E; p=0.0495, G; p=0.0002, I; p=0.7760, K; p=0.3910, M; p=0.0095) and between worker and soldier terminal larvae (B; p=0.9773, D; p=0.6789, F; p=0.6260, H; p=0.0524, J; p=0.0027, L; p=0.6565, N; p=0.0335). The sample size for relative expression values is n=6-9. (O) Allometric slopes were compared between JHE RNAi generated individuals and YFP control (p<0.0001). Furthermore, (P) head widths (p=0.0265), and (Q) scape lengths (p=0.0053) were compared between JHE RNAi and YFP control, n=20. (R) The relative expression of *JHE* using qPCR following a 48 hours JHE RNAi versus YFP RNAi treatment (p=0.0028). (S) Developmental timing frequencies compared between JHE RNAi and YFP control for cocoon formation (p=0.3899) and (T) eclosure (p=0.0056). JHE RNAi treated individuals are represented in orange and the YFP controls are in black. Representative phenotypic differences are displayed in U for YFP and *JHE* RNAi.

Furthermore, *Cad96ca* is higher in developing soldiers as compared to minors (Fig. 4*G*: p=0.0003) and trending higher in terminal soldier than minors (Fig. 4*H*; p=0.0524) while, *Fgfr1* is higher in expression in terminal soldiers as compared to minors (Fig. 4*J*; p=0.0027) and not-significantly different at the developing timepoint (Fig. 4I; p=0.7760). Finally, for degradation, we looked at the *juvenile hormone esterase* (*JHE*) and the *juvenile hormone epoxide hydrolase* (*JHEH*), two critical degraders of JH in insects (38–40). While there are multiple paralogs of *JHE* (41), we focused on the paralog that has been extensively demonstrated to regulate caste-specific minor-soldier JH-dependent behaviours in *C. floridanus* (42). We found that *JHE* and *JHEH* were significantly lower in expression in developing soldier larvae compared to minors (Fig. 4*C*; p<0.0001 and Fig. 4*E*; p=0.0495 respectively) and then are non-significantly different at the terminal timepoint (Fig. 4*D*; p=0.6789 and *F*; p=0.6260), but remain effectively shut-off compared to the early undifferentiated stage (Fig. 4*D* and *F*). Collectively, these results suggest that rather than synthesis, JH pathway activity may be regulated at the degradation and reception levels to alter ant caste determination and differentiation in *C. floridanus*.

### JH Degradation Influences Head-to-Body Allometry and Soldier Determination

It is possible that *Kr-h1* specifically, and JH activity more generally, may be higher in terminal soldier larvae due to the early decrease in JH degradation in developing soldiers that we found. In order to determine if the specific suppression of JH degradation promotes soldier-specific development, we synthesized RNAi using dsRNA of *JHE* and microinjected bipotential larvae to characterize caste-specific phenotypic effects. *JHE* RNAi significantly shifts the slope and therefore head-to body-allometry in comparison to the control, producing size variation that includes soldiers (Fig. 4*O*; p<0.0001 and Fig. 4*U*). This *JHE* RNAi treatment increased the head size (10.5% increase; Fig. 4*P*; p=0.0265), and scape length (4.7% increase, Fig. 4*Q*; p=0.0053) compared to the YFP control, by downregulating *JHE* as confirmed by significantly reduced levels of *JHE* following a 48-hour JHE RNAi treatment (Fig. 4*R*; p=0.0028). Interestingly, this change in phenotype did not significantly change the first cocoon formation (Table S4; p>0.9999) or first eclosure (Table S4; p= 0.1348). Furthermore, the treatment did not change the developmental timing of the overall cocoon formation frequency and therefore did not delay metamorphosis (Fig. 4*S*; p=0.3899), however, it did change the developmental timing of the overall eclosure frequency (Fig. 4*T*; p=0.0056). These results along with the molecular data suggest that the regulatory level of degradation could be playing a critical role in influencing the JH signalling pathway that mediates the minor-soldier caste switchpoint.

## Discussion

### Caste switchpoint: Phenotypic and Molecular levels

Based on our hormonal and molecular data, we have been able to pinpoint the JH-mediated minor-soldier switchpoint in *C. floridanus*, where larvae are most responsive to JH and subsequently differentiate into a soldier. Our data indicates that caste determination occurs during the 4^th^ larval instar when JH is capable of increasing body and head size, generating caste-specific head-to-body allometry, extending developmental timing, and inducing soldier development (Fig. 2). Furthermore, we found that JH is also capable of influencing allometry before this sensitivity period, albeit by impacting intercept rather than slope. This indicates that depending on timing, JH is capable of generating an array of allometric variation that could provide raw materials for natural selection. Although, we have determined the JH-mediated minor-soldier switchpoint, it remains unknown when the queen-worker switchpoint takes place for *C. floridanus.* In *Pheidole*, *Myrmica*, *Harpganathos, Solenopsis, Aphenogaster, Cardiocondyla, Plagiolepis, and Odontomachus* JH is known to regulate the queen-worker switchpoint (16, 21, 22, 43–46), however this has yet to be explored in *Camponotus* and needs to be further investigated. At the molecular level we found that during caste-specific differentiation, soldier-destined larvae have lower expression of the JH degraders *JHE* and *JHEH* and higher levels of expression of the JH receptor *Cad96ca* and co-factor *Taiman* relative to their minor worker-destined counterparts (Fig. 4*C*, *E*, *G* and *M*). Subsequently, by the end of caste-specific differentiation, soldier-destined larvae have higher expression of the JH receptor *Fgfr1* and maintain higher expression of *Taiman* (Fig. 3*J* and *N*). Together, this expression data across the different regulatory levels of JH signalling (Fig. 5*A*) provide insight into the roles different components of the JH signalling pathway play during the caste differentiation process. Furthermore, by taking a molecular approach we were able to more confidently pinpoint the timing of the minor-soldier switchpoint. Using *Kr-h1* as a marker, we were able to assess the sensitivity of the JH signalling pathway across a 4-day Methoprene treatment time course. We found that in response to treating 4^th^ instar larvae with methoprene, middle 4^th^ instar larvae had high levels of *Kr-h1* following a 4-day time course relative to early 4^th^ instar larvae, suggesting that the JH signalling pathway is more responsive at this precise time in larval development.

**Figure 5.**
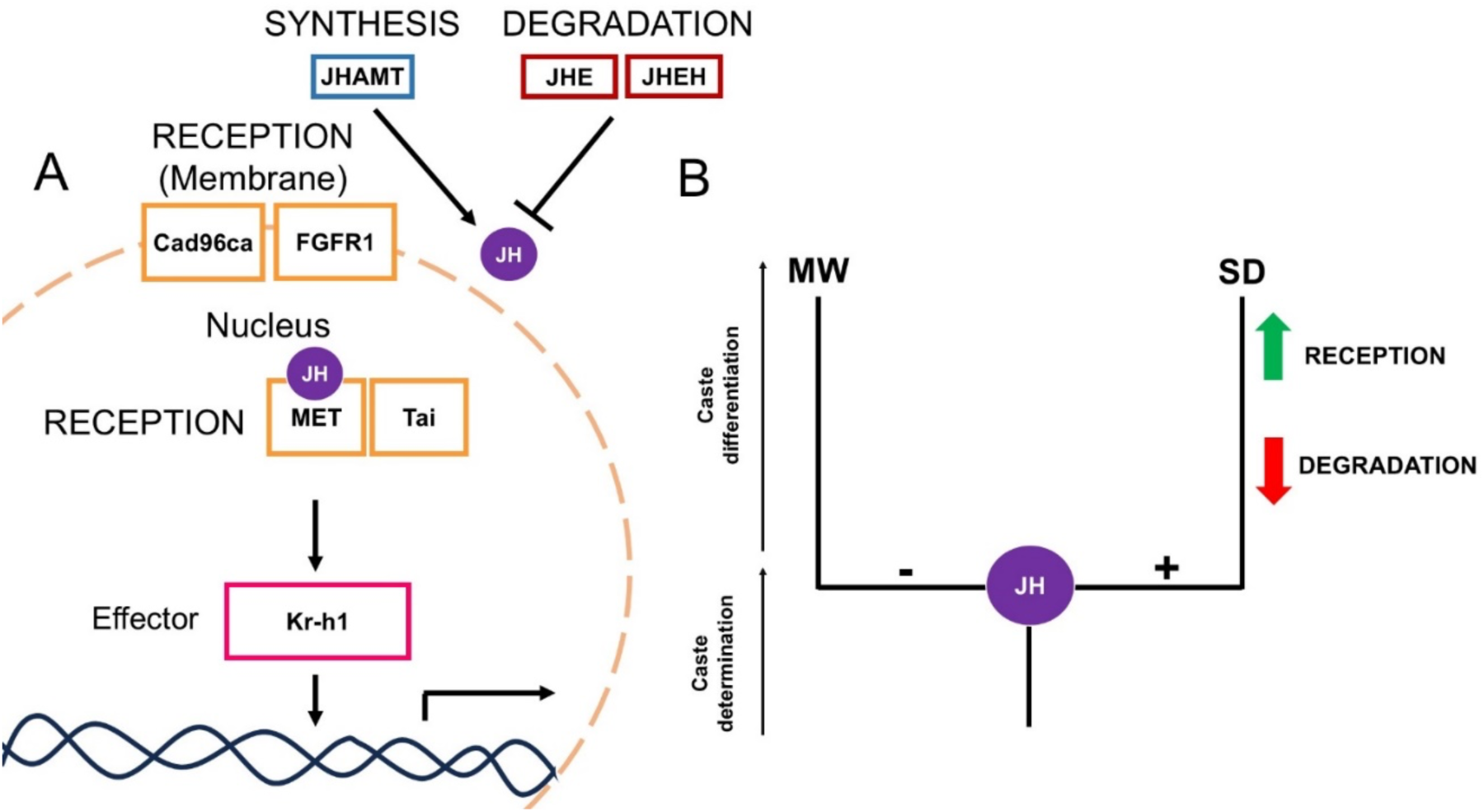
JH signalling and its differential regulation mediates the minor-soldier switchpoint in *C. floridanus*. (A) JH signalling pathway components are represented at the: (1) Synthesis (JHAMT); (2) Degradation (JHE and JHEH); and (3) Reception (membrane; Cad96ca and FGFR1, within the nucleus; MET and co-factor Tai) levels. The effector Kr-h1 is downstream of these three levels. These 3 possible levels of JH regulation that themselves can be differentially regulated in a caste-specific manner to enable caste-specific development in ants. Rather than synthesis, degradation and reception are differentially regulated to facilitate soldier development in *C. floridanus*. (B) The direction of relative expression levels for JH reception and degradation components facilitating soldier-specific development, where green arrow represents upregulation, and red arrow represents downregulation. MW is minor-worker and SD is soldier.

### Possible Endocrinological Paths to Soldier Development

Two alternative models at the endocrinological level have been suggested to lead to the evolution of the castes in ants: (1) differential sensitivities of JH through receptor distribution and abundance; (2) caste-specific differential JH levels through either regulation of synthesis or degradation (47). Based on what we have found, soldier development in *C. floridanus* is facilitated by both differential sensitivities to JH through differential receptor levels (higher) and differential levels of JH through differential degradation (lower) (Fig. 5*B*). We found the RTK membrane receptors *Cad96ca* and *Fgfr1* alongside the *Taiman* co-factor are together upregulated during soldier development, potentially functioning to increase JH signalling sensitivity. Furthermore, we found that *JHE* and *JHEH* are downregulated during soldier development and, based on our RNAi perturbation of *JHE*, the downregulation of *JHE* plays a functional role in generating larger individuals with a shift in head-to-body allometry reflective of soldiers (Fig. 4*O*). These *JHE* findings provide support that JH degradation may play a critical role in soldier development in parallel to the potential role of JH reception. Based on comparative genomics across ants in search for potential factors associated with caste dimorphism, Cad96ca, *Fgfr1*, and *JHE* were all implicated (9). Altogether, it is both through the regulation of JH levels and sensitivity that has led to the evolution of a minor-soldier switchpoint and JH-mediated soldier caste differentiation in *C. floridanus*.

In contrast to our JH treatments that extended developmental timing in a manner that would enable increased growth, our JHE RNAi did not delay metamorphosis (Fig. 4*S*) and therefore the increased growth that occurred may be through a developmental timing-independent mechanism. Similar to this, it was previously proposed that a developmental timing-independent sizing mechanism is how DNA hypomethylation increases the size of developing *C. floridanus* individuals (7). Therefore, this timing-independent mechanism regulating sizing may be regulated through the interplay of epigenetic and hormonal activity. In *C. floridanus*, an additional layer of complexity in developmental timing regulation may emerge from orally-transferred social fluids. Specifically, individuals orally transfer JH and juvenile hormone esterases (41, 48), and transmitted JH can influence developmental timing (48). Finally, rather than through the regulation of JH degradation, the developmental timing extension that occurs during soldier development in *C. floridanus* may be influenced by JH reception, which remains to be explored.

*Pheidole* and *Camponotus* are two hyperdiverse genera, both numbering over 1000 species that have diverged over 100 million years ago (23). These two groups have independently evolved soldier castes and despite their differences, seem to have similar developmental periods where JH can affect caste determination and development in a switch-like manner. It is well established that the minor-soldier caste mediated switchpoint in *Pheidole* is in the last instar (18, 47, 49), and based on our findings this switchpoint is also in the final instar for *C. floridanus*. While we characterized the differences in gene expression of the major components of the JH pathway in *C. floridanus*, this should be further explored in *Pheidole* to elucidate the evolution of developmental mechanisms underlying the independent origin of minor-soldier caste switchpoints between species.

### Contrasting developmental and adult caste-specific action of JH and JHE

JHE is differentially expressed between castes in the brain of adult *C. floridanus*, where JHE is elevated in soldiers compared to minors (26, 42). JHE downregulates JH in the brain by degrading JH at the blood-brain barrier (42) and it has been established that JH in adults promotes foraging behaviour (26, 42). Similarly, in *Apis mellifera*, increased JH levels led to an upregulation of foraging behaviours in adults (50). Conversely, we show JH promotes the development of soldiers in *C. floridanus*, which have lower *JHE* compared to minors, and that a decrease in JHE is critical for the regulation of size increase and soldier production (Fig. 2 and 4). Based on the literature and our findings it is suggested that both JH and the caste-specific function of its degradation may have opposite functions in the regulation of caste-specific development versus caste-specific behaviour in adults across social insects.

### GxE mechanisms in *C. floridanus*: Epigenetics & Hormones

Across animals, hormones and epigenetic mechanisms underlie developmental plasticity and together with genetic variation, the generation of phenotypic variation (51, 52). To date, few emerging model organisms have explored the interplay of different molecular mechanisms in mediating GxE interactions (53–57). In *C. floridanus*, recent studies have demonstrated the role of histone modifications involved in minor- and soldier-specific behaviours (26, 58, 59). In the context of *C. floridanus*, developmental epigenetic mechanisms have been demonstrated to be associated with caste-specific development and capable of regulating size variation. Specifically, DNA methylation is lower in soldiers than minor workers at the genome-wide level, these DNA methylation differences generate size variation, and DNA methylation regulates the activity of the critical cell signalling receptor *Epidermal growth factor receptor* (*Egfr*) to generate quantitative size variation (7). Here, we demonstrate that alongside caste-specific epigenetic regulation, the JH pathway plays an essential role in regulating allometric sizing, caste determination, and caste differentiation. Future research should explore other epigenetic and hormonal processes as well as address the interplay between them in the context of GxE.

## Materials and Methods

### Ant husbandry

*C. floridanus* were collected from Gainesville Florida, and colonies were kept in an incubator at 25 °C, 60% humidity, with a 12-hour light dark cycle. The ants had a constant supply of water and sugar water. They were fed three times a week with mealworms and Bhatkar & Whitcomb diet (60).

### Larval selection for manipulations and natural gene expression

*C. floridanus* larvae used for JH experiments were selected based on body length: 3^rd^ instar 2-2.7mm, early 4^th^ instar 2.8-3.4mm, and middle 4^th^ instar 3.5-4.2mm and contained a brown gut to indicate ongoing development of the larva. *C. floridanus* larvae used for expression data across caste were selected based on body length: undifferentiated 3-3.2mm, developing (based on gut color/morphology) minor 3.8-4.2mm, terminal minor 4.7-5.5mm, developing soldier (based on gut color/morphology) 6.1mm and greater, and terminal soldier 6.7mm and greater. The undifferentiated, developing minor, and developing soldier groups had to possess a brown gut to indicate ongoing larval development. Terminal minors and soldiers had to possess a black gut to indicate the end of larval development. *C. floridanus* larvae used for both temperature experiments and JHE RNAi were done with middle 4^th^ instar larvae. All larvae were either placed in the −80° freezer to be used for expression data or used immediately for experimental purposes.

### Hormonal manipulations

Larvae were topically treated with either 1ul of 5mg/ml of methoprene (Sigma-Aldrich; JH analog), or 1ul of acetone (control). Larvae were reared in a box with minor worker ants to care for them in a 1:2 larvae-adult ratio. Larvae and minor workers for each experiment were collected from the same colony. They had a constant supply of water, were fed 3 times a week with 3 mealworms cut into pieces and the Bhatkar & Whitcomb diet (60), and were either kept in an incubator with the same parameters listed above or kept at 27°C for the temperature experiment. Larva treated for expression data were collected after 24-, 48-, 72-, or 96-hour timepoints and placed into 1.5ml tubes and stored in the −80 °C freezer for RT-qPCR. Larvae treated for phenotypic measurements were collected upon eclosure and were preserved in 70% ethanol for dissection and microscopy.

### Developmental Timing

Daily observations of larvae were conducted to monitor developmental timing of cocoon formation and eclosure of ants across conditions. Adults were collected upon eclosure and stored in individual 1.5ml tubes containing 70% ethanol. Following the collection of all individuals from an experiment, the left antenna, and the head were dissected. The rest of the ant was placed back into ethanol for later use.

### Microscopy and Morphometrics

The scape and head width were measured using ZEN Pro 3.2 Software at 25x magnification using a Zeiss Axio Zoom V16 stereomicroscope. Head width was used to differentiated between minor and soldier worker castes and scape length was used as a proxy for body size. These are common parameters used for *Camponotus* species (30) and *C. floridanus* specifically (7).

### RNA extraction & cDNA synthesis

Samples were removed from the −80° freezer and placed in an ice block, then total RNA exaction and purification was performed and used a mix of the Trizol method and the Direct-zol RNA MicroPrep kit (Zymo Research). *C. floridanus* used one larva per sample. Each sample was hand homogenized in 200ul of Trizol. Following the initial homogenization, 800ul of Trizol was added and the samples were left to incubate at room temperature for 5 minutes. 200ul of chloroform was added to induce a phase separation and left to incubate for 3 minutes at room temperature. The samples were centrifuged at 12,000 ×g for 15 minutes at 4°C. The aqueous phase was extracted, placed in a new 1.5ml tube and combined with equal parts 99% ethanol (~500ul). The RNA extraction and purification proceeded using the directions of the Direct-zol RNA MicroPrep kit (Zymo Research). DNase treatment was conducted using Turbo DNase (Invitrogen). Samples were normalized at the RNA level by estimating the concentrations using a nanodrop (ThermoFisher). 500ng of cDNA was synthesized using the Super Script IV First-strand Synthesis System (Invitrogen). Each sample contained 1 larva. RNA was initially diluted to 500ng/sample. RNA and cDNA used a 1/20 dilution as the working concentration for RT-qPCR.

### RT-qPCR

Following RNA extraction and cDNA synthesis qPCR was performed to determine relative expression of key genes in the JH pathway. RT-qPCR primers were made with at least one primer across an exon-exon boundary (Table S5) and RT-qPCR was conducted with SSO advanced SYBR green (BioRad) and ran on a C1000 CFX96 machine (BioRad). The ΔΔCT (61) method was used to calculate relative fold changes. This method requires an internal normalization gene and *RpL32* (also known as rp49) was used, as it is a common gene used for normalization in *Drosophila melanogaster* (62) social insects (63) and *C. floridanus* specifically (7, 27) in the context of hormonal and epigenetic manipulations and various developmental stages. The difference in ct values between samples that had been normalized yielded the ΔΔCT values. Fold changes were calculated using 2^-ΔΔCT^ and were plotted in a bar graph.

### RNAi JHE

In order to functionally interrogate the potential role of JH degradation in soldier development, we developed a dsRNA approach that could target all JHE paralogs simultaneously. dsRNA specifically targeted the JHE paralog implicated in caste-specific behaviours in *C. floridanus* (XM123456789; 100% identity) (42), and is >86% identity to remaining JHE paralogs across the position of the gene that dsRNA was designed (Fig. S3). The *JHE* fragment was cloned using primers F: <u>GGA GAA GTG CCA GTT CGC A</u> and R: <u>GTG GCTTTA TAA AGC CTA CCG C</u> with an amplicon of 498bp, which was further verified by Sanger sequencing (Genome Quebec). This fragment was PCR amplified and incorporated T7 overhangs (non-underlined) using primers; F: TAA TAC GAC TCA CTA TAG GG<u>G GAG AAGTGC CAG TTC GCA</u> and R: TAA TAC GAC TCA CTA TAG GG<u>G TGG CTT TAT AAA GCC TACCGC</u> and checked on a gel. The PCR amplicon was purified using the QIAquick PCR purification kit (Qiagen). dsRNA was synthesized by creating 2 reactions of the following combination; 20ul of 5x transcription buffer, 1ul of RNAse out, 10ul of 10nM NTPs, 1ug of DNA, 4ul of RNA polymerase and up to 100ul of water followed by an incubation at 37°C for 4 hours. 2uL of DNAse was added and incubated for 10 minutes at 37°C. The synthesis then followed the protocol outlined in the Megaclear kit from (ThermoFisher). Finally, the 2 reactions were combined and concentrated by adding 12ul of 5M NaCl, and 600ul of ice cold 95% ethanol, followed by a 30 minute incubation on ice. The reaction was centrifuged at 4°C for 10 minutes at 17,000 ×g and supernatant was discarded. 600ul of ice cold 70% ethanol was added and centrifuged at 17,000 ×g for 5 minutes at 4°C. Supernatant was discarded and the pellet was dried at 37°C for 5-10 minutes. It was then re-suspended in 20ul of injection buffer and the concentration was estimated by nanodrop. The JHE synthesized dsRNA and YFP control were injected at the same concentration, approximately 4000ng/ul. With the aid of a Zeiss SteREO Discovery V8 microscope, the dsRNA solution was microinjected into bipotential larvae using an Eppendorf CellTram 4r (oil) microinjector, controlled by a Narishige micromanipulator and reared in a similar manner to the JH treatments described above. Microinjection needles were made using a Sutter Instrument P-97 needle puller.

### Statistical analysis

All statistical analysis was performed using Graphpad Prism. RT-qPCR fold changes were graphed in bar graphs and one-way ANOVAs were run for natural expression levels across developmental stages; unpaired two-tailed t-test were conducted for hormonal treatments to determine significance between control and treatment groups with a p-value of <0.05. JHE RNAi treatments used an unpaired one-tailed t-test to test the efficacy of JHE RNAi knockdown in decreasing the expression level of JHE. All statistical tests for RT-qPCR analysis used a p-value <0.05.

Phenotypic measurements were plotted on a XY scatter plot and statistically analyzed using GraphPad prism. A simple linear regression was used to determine caste-specific differences between treatment and control groups. Unpaired two-tailed T-tests were conducted to determine head width and scape length differences between control and treatment groups with a p-value of <0.05 for all natural and methoprene treatments. An unpaired one-tailed t-test was conducted to test if JHE RNAi would lead to the predicted increase in body size and head size (based on the natural expression across developmental stages).

Developmental timing extensions were calculated by averaging the first pupation or eclosure from each trial and using the following equation to determine percent extension. Percent extension = ((treatment – control)/control) x 100%. Furthermore, the statistical significance of developmental differences was assessed using an unpaired two-tailed t-test with a p-value <0.05. The significance of pupation and eclosure frequencies were calculated using unpaired two-tailed t-tests with a p-value <0.05.

## Acknowledgments

We thank L. Davis and the Abouheif lab for help with ant collection. We also thank A. Rajakumar for comments and Rajakumar Lab members for both comments and husbandry support and the Abouheif lab for experimental advice. We acknowledge the support of the Natural Sciences and Engineering Research Council of Canada (NSERC) [RGPAS-2021-00006; RGPIN-2021-04399; RTI-2021-00710]; Canadian Foundation for Innovation and the Ontario Research Fund [40142].

## Supplemental Information

**Fig. S1.**
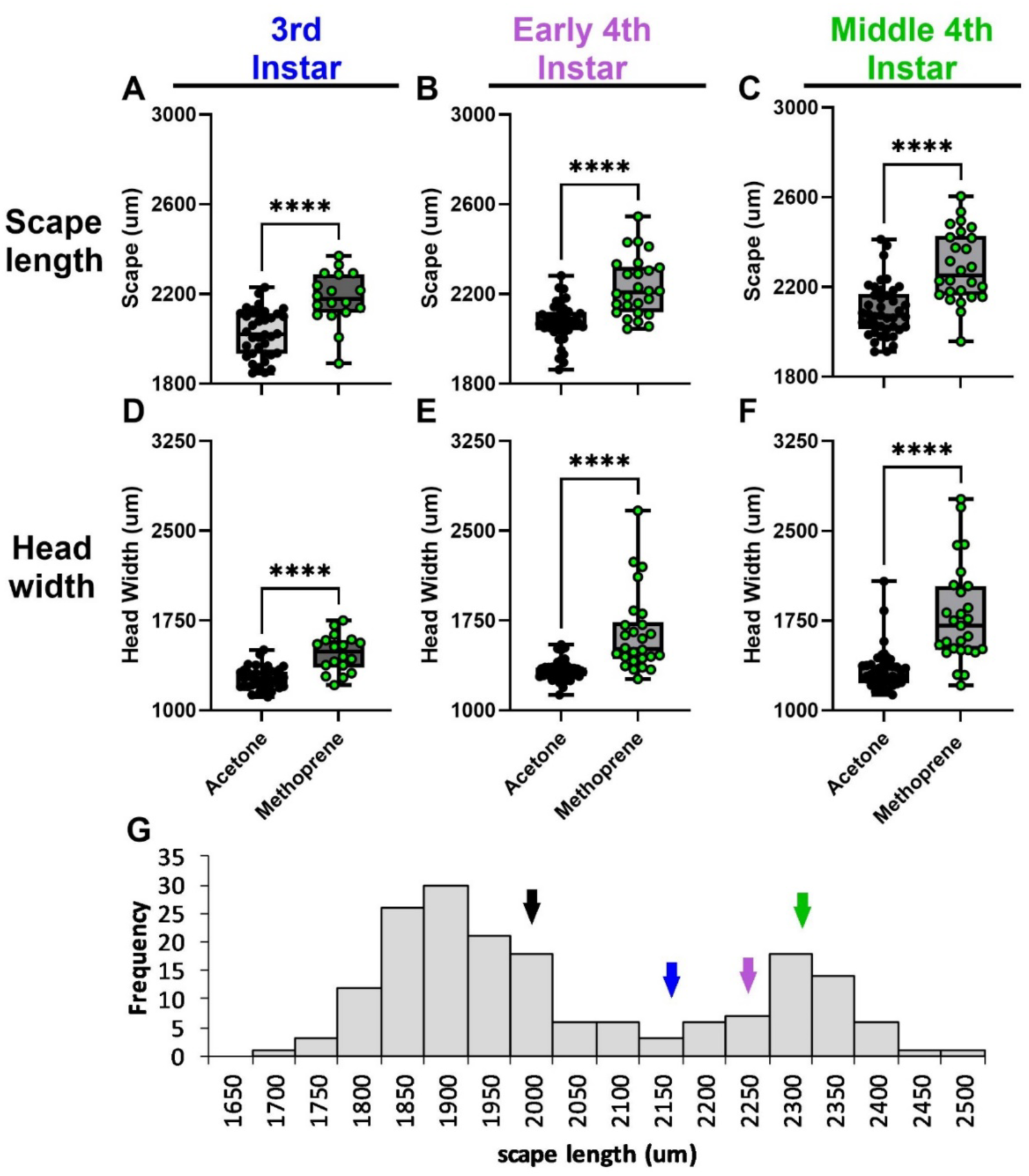
Changes in scape lengths and head widths of adult ants following JH treatment at different developmental stages. (A-C) Scape lengths of individuals generated after methoprene treatment of 5mg/ml when treated at: (A) 3^rd^ instar (p<0.0001), (B) early 4^th^ instar (p=0.0008), and (C) middle 4^th^ instar treatments (p<0.0001) were compared between methoprene treatment and acetone control. (D-F) Head widths of individuals generated after methoprene treatment when treated at: (D) 3^rd^ instar (p=<0.0001), (E) early 4^th^ instar (p=0.0006), and (F) middle 4^th^ instar (p=<0.0001) were compared between methoprene treatment and acetone control. Methoprene treated individuals are represented in green and acetone control is represented in black. (G) Arrows representing the control (black), and methoprene treatments: 3^rd^ instar (blue), early 4^th^ instar (purple), and middle 4^th^ instar (green) demonstrate the scape lengths of treated samples in comparison to scape measurements across a mature *C. floridanus* colony adapted from that previously described (7).

**Fig. S2.**
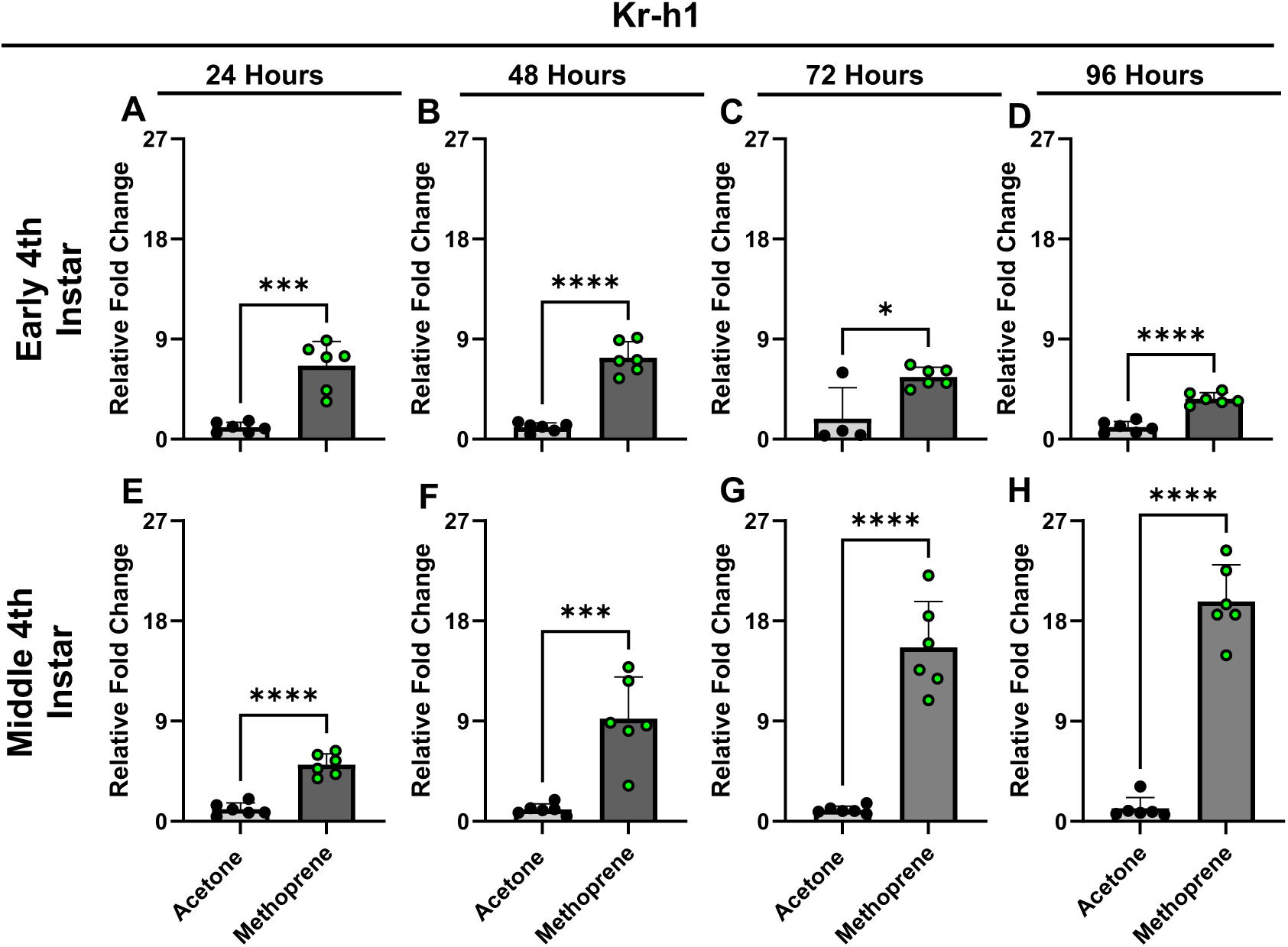
*Kr-h1* expression following a methoprene treatment time-course during larval stages. Relative expression of *Kr-h1* during early 4^th^ instar following a methoprene treatment at: (A) 24-hours (p=0.0001), (B) 48-hours (p<0.0001), (C)72-hours, and (D) 96-hours (p<0.0001) was compared between methoprene treatment and acetone control. Relative expression of *Kr-h1* during middle 4^th^ instar following a methoprene treatment at: (E) 24-hours (p<0.0001), (F) 48-hours (p=0.0004), (G)72-hours (p<0.0001), and (H)96-hours (p<0.0001) was compared between methoprene treatment and acetone control. Methoprene treated individuals are represented in green and acetone control is represented in black, n=6.

**Fig. S3.**
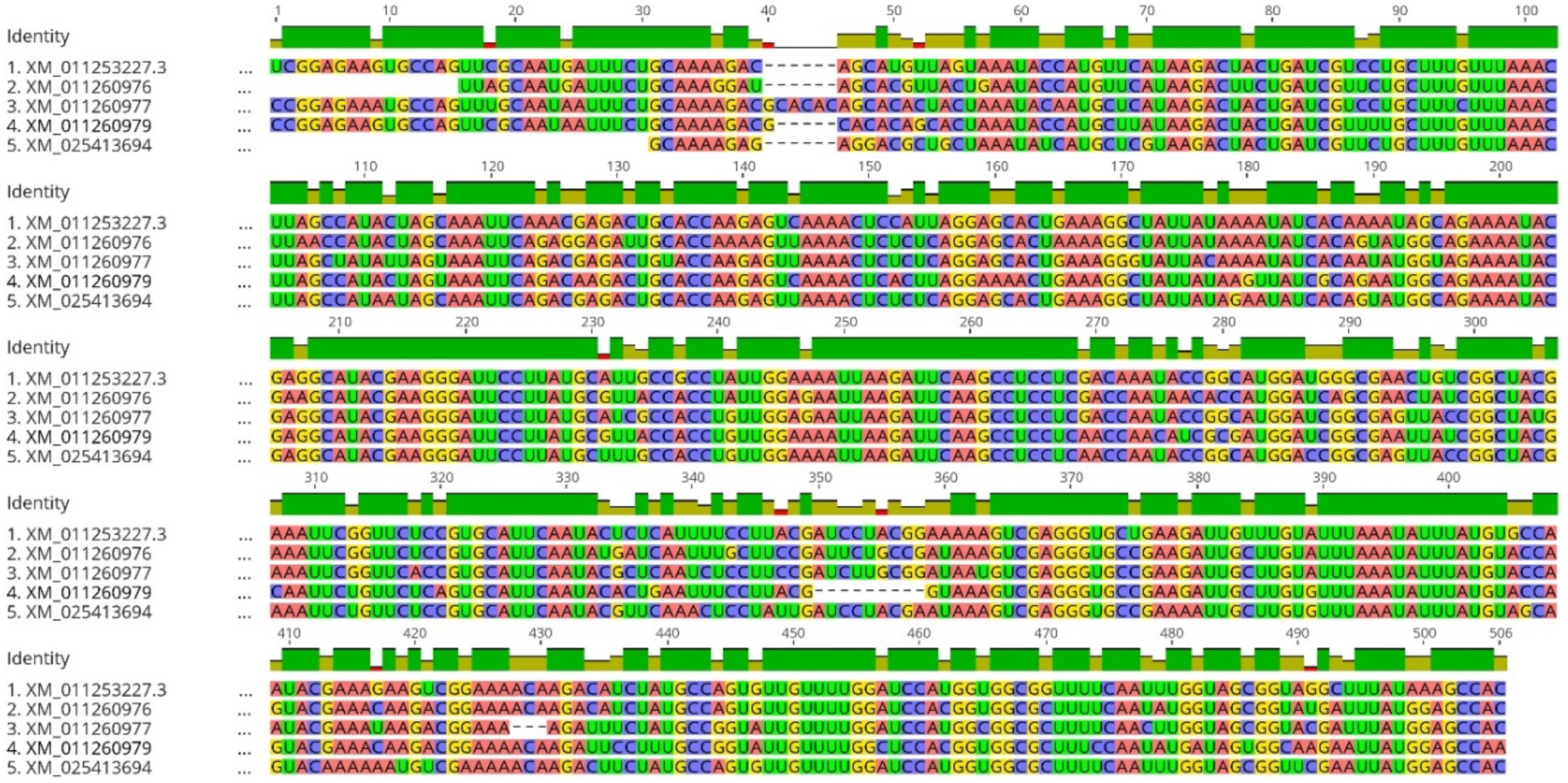
Sequence alignment of the RNAi targeted region for *C. floridanus* JHEs. Indicated are the 5 JHE paralogs with the NCBI accession numbers indicated on the left. The dsRNA fragment is designed to specifically target (100% identical) XM_011253227.3, which is the JHE known to specifically mediate *C. floridanus* minor-soldier caste-specific behaviours (42). This fragment has 86% pairwise identity with the other JHE paralogs, enabling the targeting of all JHEs facilitating the suppression of JH degradation more broadly to address the role of JH degradation in soldier-specific development.

**Table S1.** Difference in timing to first pupation between methoprene and acetone treatments across larval stages. The first pupation was recorded as the day when the first individual from each condition became a cocoon. The first pupation across trials was averaged and the percent change was calculated using (average methoprene – average acetone)/average acetone x 100%.

| First pupation |  |  |  |  |  |  |  |  |
| --- | --- | --- | --- | --- | --- | --- | --- | --- |
| Trial | 3rd Instar |  | Trial | Early 4th Instar |  | Trial | Middle 4th Instar |  |
|  | Acetone | Methoprene |  | Acetone | Methoprene |  | Acetone | Methoprene |
| 6 | 11 | 15 | 5 | 12 | 16 | 1 | 6 | 25 |
| 8 | 23 | 19 | 7 | 10 | 23 | 2 | 7 | 20 |
| 10 | 10 | 17 | 9 | 9 | 16 | 4 | 8 | 16 |
| 13 | 11 | 23 | 16 | 10 | 19 | 11 | 7 | 21 |
| 15 | - | - | 19 | 12 | 21 | 17 | 6 | 10 |
| 18 | 12 | 36 | 22 | 12 |  | 20 | 9 | 18 |
| 21 | 18 | 24 | 26 | 12 | 16 | 23 | 6 | 18 |
| 24 | 12 | 24 |  |  |  |  |  |  |
| 25 | 11 | 16 |  |  |  |  |  |  |
| Average | 13.50 | 21.75 | Average | 11 | 18.5 | Average | 7 | 18.28571429 |
| Difference in days |  | 8.25 | Difference in days |  | 7.5 | Difference in days |  | 11.28571429 |
| % Change |  | 61.11111111 | % Change |  | 68.18181818 | % Change |  | 161.2244898 |
| Conclusion | 61% delay in first pupation relative to acetone |  | Conclusion | Methoprene has a 68% delay in first pupation relative to acetone |  | Conclusion | Methoprene has a 161% delay in first pupation relative to acetone |  |
| P-value | 0.0129 |  | P-value | <0.0001 |  | P-value | <0.0001 |  |

**Table S2.** Difference in timing to first eclosure between methoprene and acetone treatments across larval sizes. The first eclosure was recorded as the day when the first individual from each condition eclosed from their cocoon and emerged as an adult. The first eclosure across trials was averaged and the percent change was calculated using (average methoprene – average acetone)/average acetone x 100%.

| 3rd Instar |  |  | Early 4th Instar |  |  | Middle 4th Instar |  |  |
| --- | --- | --- | --- | --- | --- | --- | --- | --- |
| Trial | Acetone | Methoprene | Trial | Acetone | Methoprene | Trial | Acetone | Methoprene |
| 6 | 37 | 40 | 5 | 37 | 43 | 1 | 29 | 52 |
| 8 | 50 | 45 | 7 | 34 | 50 | 2 | 32 | 49 |
| 10 | 34 | 46 | 9 | 33 | 43 | 4 | 32 | 42 |
| 13 | 38 | 52 | 16 | 35 | 45 | 11 | 31 | 51 |
| 15 | - | - | 19 | 36 | 51 | 17 | 31 | 45 |
| 18 | 36 | - | 22 | 33 |  | 20 | 33 | 50 |
| 21 | 36 | 36 | 26 | 30 | 38 | 23 | 30 | 40 |
| 24 | 33 | - |  |  |  |  |  |  |
| 25 | 32 | 39 |  |  |  |  |  |  |
| Average | 37 | 43 | Average | 34 | 45 | Average | 31.14285714 | 47 |
| Difference in days | 6 |  | Difference in days | 11 |  | Difference in days | 15.85714286 |  |
| % Change | 16.21621622 |  | % Change | 32.35294118 |  | % Change | 50.91743119 |  |
| Conclusion | Methoprene has a 16% delay in eclosure relative to acetone |  | Conclusion | Methoprene has a 32% delay in eclosure relative to acetone |  | Conclusion | Methoprene has a 51% delay in eclosure relative to acetone |  |
| P-value | 0.0751 |  | P-value | 0.0002 |  | P-value | <0.0001 |  |

**Table S3.**
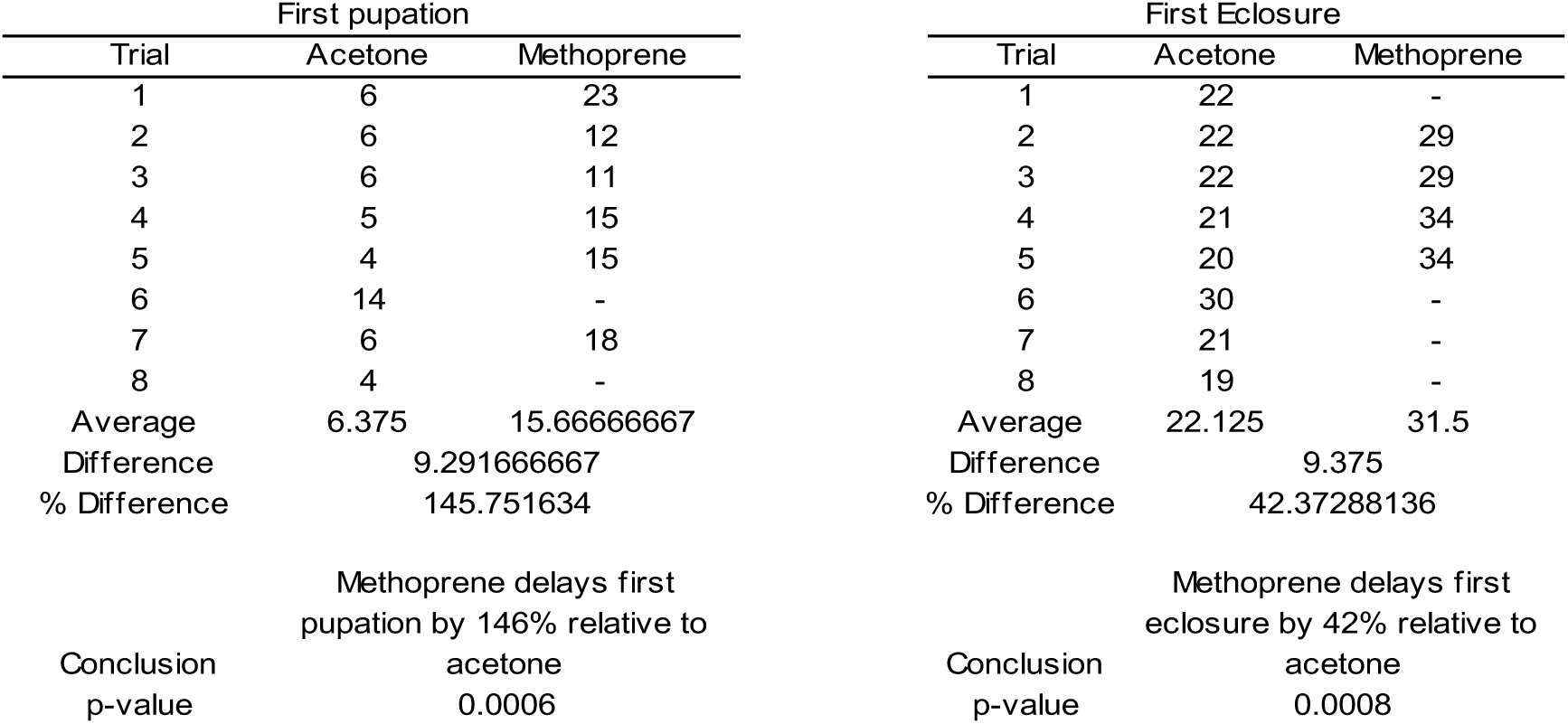
Changes in developmental timing of first pupation and eclosure after methoprene treatment reared at 27°C. The first pupation was recorded as the day when the first individual from each condition became a cocoon. The first eclosure was recorded as the day when the first individual eclosed from their cocoon and emerged as an adult. The data was averaged and the percent change was calculated using (average methoprene – average acetone)/average acetone x 100%.

**Table S4.** Changes in developmental timing of first pupation and eclosure after JHE RNAi treatment. The first pupation was recorded as the day when the first individual from each condition became a cocoon. The first eclosure was recorded as the day when the first individual eclosed from their cocoon and emerged as an adult. The data was averaged and the percent change was calculated using (average methoprene – average acetone)/average acetone x 100%.

| First pupation |  |  | First Eclosure |  |  |
| --- | --- | --- | --- | --- | --- |
| Trial | Yfp | JHE RNAi | Trial | Yfp | JHE RNAi |
| 12 | 7 | 9 | 12 | 28 | 29 |
| 13 | 8 | - | 13 | 28 | - |
| 14 | 7 | 10 | 14 | 33 | 33 |
| 15 | 9 | 7 | 15 | 32 | 35 |
| 16 | - | 7 | 16 | - | 33 |
| 17 | 4 | 15 | 17 | 25 |  |
| 18 | 16 | 16 | 18 | - | - |
| 19 | 21 | 7 | 19 | - | - |
| 20 | 6 | 7 | 20 | 31 | 31 |
| Average | 9.75 | 9.75 | Average | 29.5 | 32.2 |
| Difference |  | 0 | Difference |  | 2.7 |
| % Difference |  | 0 | % Difference |  | 9.152542373 |
| Conclusion | No difference in first pupation for Yfp and JHE RNAi |  | Conclusion | JHE RNAi delayed first eclosure by 9.2% |  |
| P-value | >0.9999 |  | P-value | 0.1348 |  |

**Table S5.**
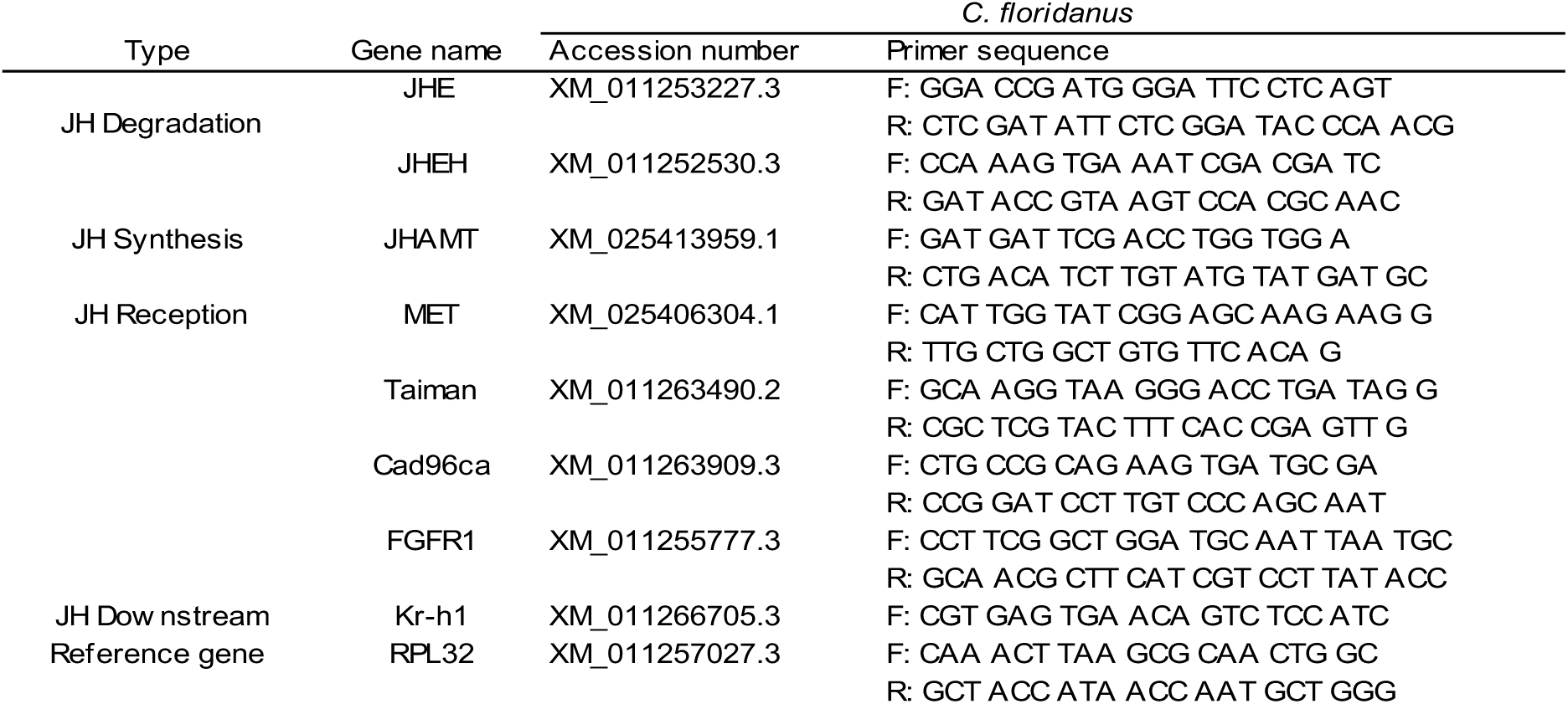
Primers used for qPCR to determine relative gene expression. Primer sequences for all JH pathway components targeted as well as the accession number from NCBI used to design the primers. At least one primer for each gene spanned an exon-exon junction to prevent intronic amplification.

## References

1. B. Hölldobler, E. O. Wilson, The ants (Harvard University Press., 1990).

2. E. O. Wilson, Causes of Ecological Success: The Case of the Ants. The Journal of Animal Ecology 56, 1 (1987).

3. B. Holldobler, E. O. Wilson, The superorganism: the beauty elegance and strangeness of insect societies (WW Norton & Company, 2009).

4. D. E. Wheeler, The Developmental Basis of Worker Caste Polymorphism in Ants. The American Naturalist 138, 1218–1238 (1991).

5. R. Rajakumar, et al., Social regulation of a rudimentary organ generates complex worker-caste systems in ants. Nature 562, 574–577 (2018).

6. E. O. Wilson, The Origin and Evolution of Polymorphism in Ants. The Quarterly Review of Biology 28, 136–156 (1953).

7. S. Alvarado, R. Rajakumar, E. Abouheif, M. Szyf, Epigenetic variation in the Egfr gene generates quantitative variation in a complex trait in ants. Nat Commun 6, 6513 (2015).

8. J. Gadau, J. Heinze, B. Hölldobler, M. Schmid, Population and colony structure of the carpenter ant *Camponotus floridanus*. Molecular Ecology 5, 785–792 (1996).

9. J. Vizueta, et al., Adaptive radiation and social evolution of the ants. Cell S0092867425006178 (2025). 10.1016/j.cell.2025.05.030.

10. M. J. West-Eberhard, Phenotypic Plasticity and the Origins of Diversity. Annual Review of Ecology and Systematics 20, 249–278 (1989).

11. M. J. West-Eberhard, Developmental plasticity and evolution. (Oxford University Press., 2003).

12. L. Passera, Différenciation des soldats chez la FourmiPheidole pallidula Nyl. (Formicidae Myrmicinae). Ins. Soc 21, 71–86 (1974).

13. W. Goetsch, Die Staaten der Ameisen (Springer Berlin Heidelberg, 1937).

14. F. Smith, Effect of Reduced Food Supply upon the Stature of Camponotus Ants. Entomological News 53, 133–135 (1942).

15. F. Smith, Nutritional Requirements of Camponotus Ants1. Annals of the Entomological Society of America 37, 401–408 (1944).

16. L. Passera, J. P. Suzzoni, Le role de la reine dePheidole pallidula (Nyl.) (Hymenoptera, Formicidae) dans la sexualisation du couvain après traitement par l’hormone juvénile. Ins. Soc 26, 343–353 (1979).

17. D. E. Wheeler, H. Frederik Nijhout, Soldier determination in Pheidole bicarinata: Effect of methoprene on caste and size within castes. Journal of Insect Physiology 29, 847–854 (1983).

18. D. E. Wheeler, H. F. Nijhout, Soldier Determination in Ants: New Role for Juvenile Hormone. Science 213, 361–363 (1981).

19. H. F. Nijhout, D. E. Wheeler, Juvenile Hormone and the Physiological Basis of Insect Polymorphisms. The Quarterly Review of Biology 57, 109–133 (1982).

20. D. E. Wheeler, Developmental and Physiological Determinants of Caste in Social Hymenoptera: Evolutionary Implications. The American Naturalist 128, 13–34 (1986).

21. M. V. Brian, Caste differentiation in Myrmica rubra: The rôle of hormones. Journal of Insect Physiology 20, 1351–1365 (1974).

22. C. A. Penick, S. S. Prager, J. Liebig, Juvenile hormone induces queen development in late-stage larvae of the ant Harpegnathos saltator. Journal of Insect Physiology 58, 1643–1649 (2012).

23. J. Romiguier, et al., Ant phylogenomics reveals a natural selection hotspot preceding the origin of complex eusociality. Current Biology 32, 2942–2947.e4 (2022).

24. R. Bonasio, et al., Genomic Comparison of the Ants *Camponotus floridanus* and *Harpegnathos saltator*. Science 329, 1068–1071 (2010).

25. R. Bonasio, et al., Genome-wide and Caste-Specific DNA Methylomes of the Ants Camponotus floridanus and Harpegnathos saltator. Current Biology 22, 1755–1764 (2012).

26. K. M. Glastad, et al., Epigenetic Regulator CoREST Controls Social Behavior in Ants. Molecular Cell 77, 338–351.e6 (2020).

27. J. Gospocic, et al., Kr-h1 maintains distinct caste-specific neurotranscriptomes in response to socially regulated hormones. Cell 184, 5807–5823.e14 (2021).

28. A. C. LeBoeuf, et al., Oral transfer of chemical cues, growth proteins and hormones in social insects. eLife 5, e20375 (2016).

29. E. J. Shields, L. Sheng, A. K. Weiner, B. A. Garcia, R. Bonasio, High-Quality Genome Assemblies Reveal Long Non-coding RNAs Expressed in Ant Brains. Cell Reports 23, 3078–3090 (2018).

30. J. A. F. Diniz-Filho, C. J. Von Zuben, H. G. Fowler, M. N. Schlindwein, O. C. Bueno, Multivariate morphometrics and allometry in a polymorphic ant. Ins. Soc 41, 153–163 (1994).

31. Q. He, Y. Zhang, Kr-h1, a Cornerstone Gene in Insect Life History. Front. Physiol. 13, 905441 (2022).

32. J. Lozano, X. Belles, Conserved repressive function of Krüppel homolog 1 on insect metamorphosis in hemimetabolous and holometabolous species. Sci Rep 1, 163 (2011).

33. T. Shinoda, K. Itoyama, Juvenile hormone acid methyltransferase: A key regulatory enzyme for insect metamorphosis. Proc. Natl. Acad. Sci. U.S.A. 100, 11986–11991 (2003).

34. M. Jindra, M. Uhlirova, J.-P. Charles, V. Smykal, R. J. Hill, Genetic Evidence for Function of the bHLH-PAS Protein Gce/Met As a Juvenile Hormone Receptor. PLoS Genet 11, e1005394 (2015).

35. B. Konopova, M. Jindra, Juvenile hormone resistance gene *Methoprene-tolerant* controls entry into metamorphosis in the beetle *Tribolium castaneum*. Proc. Natl. Acad. Sci. U.S.A. 104, 10488–10493 (2007).

36. J. Lozano, T. Kayukawa, T. Shinoda, X. Belles, A Role for Taiman in Insect Metamorphosis. PLoS Genet 10, e1004769 (2014).

37. Y.-X. Li, et al., Receptor tyrosine kinases CAD96CA and FGFR1 function as the cell membrane receptors of insect juvenile hormone. eLife 13 (2025).

38. S. Debernard, et al., Expression and characterization of the recombinant juvenile hormone epoxide hydrolase (JHEH) from Manduca sexta. Insect Biochemistry and Molecular Biology 28, 409–419 (1998).

39. S. G. Kamita, et al., Juvenile hormone (JH) esterase: why are you so JH specific? Insect Biochemistry and Molecular Biology 33, 1261–1273 (2003).

40. L. L. Sanburg, K. J. Kramer, F. J. Kezdy, J. H. Law, H. Oberlander, Role of juvenile hormone esterases and carrier proteins in insect development. Nature 253, 266–267 (1975).

41. A. C. LeBoeuf, et al., Molecular evolution of juvenile hormone esterase-like proteins in a socially exchanged fluid. Sci Rep 8, 17830 (2018).

42. L. Ju, et al., Hormonal gatekeeping via the blood-brain barrier governs caste-specific behavior in ants. Cell 186, 4289–4309.e23 (2023).

43. S. Bradleigh Vinson, R. Robeau, Insect Growth Regulator Effects on Colonies of the Imported Fire Ant12. Journal of Economic Entomology 67, 584–587 (1974).

44. P. Colombel, Biologie d’Odontomachus haematodes L. (Hym. Form.) determinisme de la caste femelle. Ins. Soc 25, 141–151 (1978).

45. Ledoux, Dargagnon, formation des castes chez la fourmi Aphaenogaster senilis Mayr. Acad Sci Paris CR Ser D. (1973).

46. A. Schrempf, J. Heinze, Proximate mechanisms of male morph determination in the ant *Cardiocondyla obscurior*. Evolution and Development 8, 266–272 (2006).

47. R. Rajakumar, et al., Ancestral Developmental Potential Facilitates Parallel Evolution in Ants. Science 335, 79–82 (2012).

48. M. A. Negroni, A. C. LeBoeuf, Social administration of juvenile hormone to larvae increases body size and nutritional needs for pupation. R. Soc. open sci. 10, 231471 (2023).

49. E. Abouheif, G. A. Wray, Evolution of the Gene Network Underlying Wing Polyphenism in Ants. Science 297, 249–252 (2002).

50. G. E. Robinson, C. Strambi, A. Strambi, M. F. Feldlaufer, Comparison of juvenile hormone and ecdysteroid haemolymph titres in adult worker and queen honey bees (Apis mellifera). Journal of Insect Physiology 37, 929–935 (1991).

51. E. Abouheif, et al., “Eco-Evo-Devo: The Time Has Come” in Ecological Genomics, Advances in Experimental Medicine and Biology., C. R. Landry, N. Aubin-Horth, Eds. (Springer Netherlands, 2014), pp. 107–125.

52. S. F. Gilbert, T. C. G. Bosch, C. Ledón-Rettig, Eco-Evo-Devo: developmental symbiosis and developmental plasticity as evolutionary agents. Nat Rev Genet 16, 611–622 (2015).

53. P. M. Brakefield, et al., Development, plasticity and evolution of butterfly eyespot patterns. Nature 384, 236–242 (1996).

54. D. J. Emlen, I. A. Warren, A. Johns, I. Dworkin, L. C. Lavine, A Mechanism of Extreme Growth and Reliable Signaling in Sexually Selected Ornaments and Weapons. Science 337, 860–864 (2012).

55. C. Ge, et al., The histone demethylase KDM6B regulates temperature-dependent sex determination in a turtle species. Science 360, 645–648 (2018).

56. T. Kijimoto, A. P. Moczek, J. Andrews, Diversification of *doublesex* function underlies morph-, sex-, and species-specific development of beetle horns. Proc. Natl. Acad. Sci. U.S.A. 109, 20526–20531 (2012).

57. E. J. Ragsdale, M. R. Müller, C. Rödelsperger, R. J. Sommer, A Developmental Switch Coupled to the Evolution of Plasticity Acts through a Sulfatase. Cell 155, 922–933 (2013).

58. K. M. Glastad, L. Ju, S. L. Berger, Tramtrack acts during late pupal development to direct ant caste identity. PLoS Genet 17, e1009801 (2021).

59. D. F. Simola, et al., Epigenetic (re)programming of caste-specific behavior in the ant *Camponotus floridanus*. Science 351, aac6633 (2016).

60. A. Bhatkar, W. H. Whitcomb, Artificial Diet for Rearing Various Species of Ants. The Florida Entomologist 53, 229 (1970).

61. K. J. Livak, T. D. Schmittgen, Analysis of Relative Gene Expression Data Using Real-Time Quantitative PCR and the 2−ΔΔCT Method. Methods 25, 402–408 (2001).

62. J. Colombani, D. S. Andersen, P. Léopold, Secreted Peptide Dilp8 Coordinates *Drosophila* Tissue Growth with Developmental Timing. Science 336, 582–585 (2012).

63. T. S. Depintor, F. C. P. Freitas, N. Hernandes, F. M. F. Nunes, Z. L. P. Simões, Interactions of juvenile hormone, 20-hydroxyecdysone, developmental genes, and miRNAs during pupal development in Apis mellifera. Sci Rep 15, 10354 (2025).

